# Plant NLR targets P-type ATPase for executing plasma membrane depolarization leading to calcium influx and cell death

**DOI:** 10.1101/2020.08.30.274688

**Authors:** Hye-Young Lee, Ye-Eun Seo, Joo Hyun Lee, So Eui Lee, Soohyun Oh, Jihyun Kim, Seungmee Jung, Haeun Kim, Hyojeong Park, Sejun Kim, Hyunggon Mang, Doil Choi

## Abstract

Hypersensitive response (HR) is a robust immune response mediated by plant nucleotide-binding and leucine-rich repeat receptor (NLR). However, the early molecular event linking NLR to cell death is obscure. Here we demonstrate that NLR targets plasma membrane H^+^-ATPases (PMA) generating electrochemical potential across the membrane. CC^A^309, an autoactive N-terminal domain of pepper coiled-coil NLR (CNL), associates with PMAs and its autoactivity is affected by silencing or overexpression of PMA. CC^A^309-induced extracellular alkalization accompanied with membrane depolarization is followed by calcium influx and cell death. CC^A^309 interacts with C-terminal regulatory domain of PMA and 14-3-3 negatively affects CC^A^309-induced cell death. Moreover, pharmacological experiments with fusicoccin, an irreversible PMA activator, confirmed that CC- and CNL-mediated cell death occurred through inhibiting PMA. We propose PMAs as the primary target of plasma membrane-associated CNL to disrupt electrochemical homeostasis leading to HR cell death.

## Main text

Plants rely on the innate immune system against pathogen challenges. The immune responses are provoked by the perception of pathogen infection via the cytosolic immune receptors, designated nucleotide-binding and leucine-rich repeat receptor (NLR)^1^. Plant NLRs are subdivided into two major classes based on their N-terminal domain; coiled-coil NLR (CNL) and Toll interleukin-1 receptor NLR (TNL)^2^. The NLRs are activated upon the recognition of pathogenic effectors, resulting in defense responses including hypersensitive response (HR), a form of programmed cell death^3^. Although HR is considered as one of the most robust defense responses and facilitates restriction of pathogen growth in the infected cells, the underlying mechanisms are elusive.

Recent remarkable discoveries shed light on the molecular events of NLR-mediated cell death. Like bacterial and mammalian TIR domains, plant TIR domain in TNL cleaves NAD^+^, leading to cell death signal transduction^4,5^. Moreover, structural and biochemical characterization of *Arabidopsis* ZAR1 revealed that the N-terminal α-helix of CC domain in the activated resistosome forms a funnel-shaped structure, which is essential for ZAR1-mediated cell death^6^. However, the early molecular events in the onset of cell death by activated NLR remain to be elucidated.

Previously, we reported an Autonomous and Ancient NLR group (ANL) comprising CC domains conferring HR-mimic cell death in seed plants^7^. To take experimental advantages of small size and simple structure, CC^A^309, an autoactive CC domain in pepper ANL, was adopted to identify novel immune components underlying NLR-mediated cell death.

We identified CC^A^309 associated proteins in *Nicotiana benthamiana* by *in planta* co-immunoprecipitation (co-IP) followed by liquid chromatography-tandem mass spectrometry (Extended Data Table 1). Since CC^A^309 is localized on plasma membrane (PM), plasma membrane H^+^-ATPase3 (PMA3) among the candidates was selected for further experiments. PMAs are active proton pumps to establish the electrochemical gradient across the PM which is essential not only for the exchange of ions and solutes across the PM but also for cell viability as shown by genetic analysis in *Arabidopsis* and yeast^8,9^. Co-IP and bimolecular fluorescence complementation (BiFC) assays validated that PMA3 associates with CC^A^309 but not NbRIN4, a homolog of PM-localized *Arabidopsis* RPM1-interacting protein 4, indicating that PMA3 may specifically interact with CC^A^309 (Fig. 1a, b, Extended Data Fig. 1). The CC^A^309-induced cell death but not Bax encoding human BCL2-associated X was significantly enhanced in PMA3-silenced plants (Fig. 1c, Extended Data Fig. 2) and compromised by transient expression *PMA3* in *N. benthamiana* (Fig. 1d). These results suggest that PMA3 plays an important role in CC^A^309-induced cell death. Plants possess multiple isoforms of differentially expressed PMAs with functional redundancy^8^. We identified additional 12 putative PMAs from *N. benthamiana* genome^10^ and further revealed that PMA1 in the same phylogenetic clade of PMA3 and PMA4 in another clade also interacted with CC^A^309, suggesting that CC^A^309 associates with multiple PMAs *in planta* (Extended Data Fig. 3, 4). Consistent with these data, multiple *PMA*-silenced plants (*PMA1/2/*3) showed significantly enhanced cell death caused by CC^A^309, note that *PMA1/2/3/4*-silenced plants were not subjected to the test because of extremely abnormal growth (Extended Data Fig. 5).

**Fig. 1.**
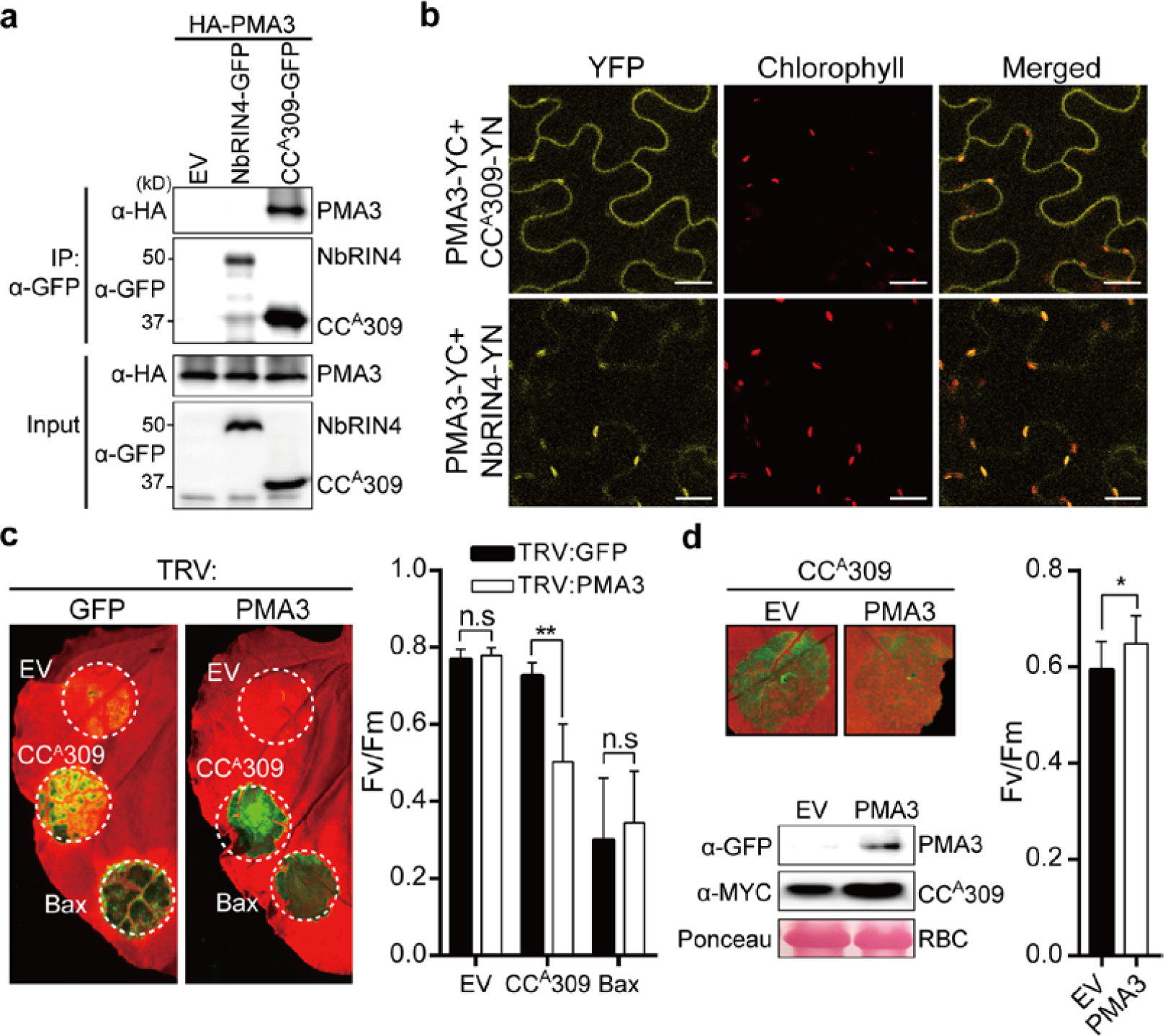
The CC^A^309-induced cell death is negatively regulated by PMA3. **a, b**, CC^A^309 not NbRIN4 associated with PMA3. *35S:HA-PMA3* was co-expressed with *35S:NbRIN4-GFP, 35S:CC*^*A*^*309-GFP* or empty vector (EV) in *N. benthamiana*. Protein extracts were immunoprecipitated with GFP antibody (IP: α-GFP) and immunoblotted with α-GFP or α-HA. Protein inputs are shown with immunoblotting before immunoprecipitation. **a**, BiFC assay in *N. benthamiana. 35S:PMA3-YC* was co-expressed with *35S:CC*^*A*^*309-YN* or *35S:NbRIN4-YN*. **b**, The YFP signals were detected by a confocal microscope after 40 hours post agro-infiltration (hpi). YFP, yellow fluorescent protein signal; Chlorophyll, autofluorescence signal; Merged, YFP merged with Chlorophyll. Scale Bar = 20 μm. NbRIN4 localized in PM is used as a negative control. **c**, Enhanced CC^A^309-induced cell death in *PMA3*-silenced plants. Silenced plants by *TRV:GFP* or *TRV:PMA3* were agro-infiltrated with CC^A^309 at three weeks after silencing. Leaves were photographed at 3-days post infiltration (dpi) (left panel) and the degree of cell death is depicted by quantification of quantum yield (Fv/Fm) (right panel). Human Bax and EV were used as positive and negative control for cell death, respectively. Significance was determined using t-test, with asterisks denoting statistically significant differences. **P < 0.01. n.s.; no significant. **d**, Decreased CC^A^309-induced cell death by overexpressing PMA3. *35S:CC*^*A*^*309-MYC* was co-expressed *35S:PMA3-GFP* or EV in *N. benthamiana*. Cell death was photographed at 5 dpi (left top) and quantified as Fv/Fm (right). The proteins were detected by immunoblotting with α-GFP or α-MYC. Ponceau S staining of RuBisCO (RBC) is loading control (left bottom). Data are represented as mean ± SD (N = 9) by triplicated independent experiments. Asterisk denotes a significant difference (t-test, *P < 0.05).

As PMA is a major H^+^ pump in the plasma membrane, we monitored the pH change in the apoplast and cytoplasm during CC^A^309-induced cell death. To measure pH values inside and outside of the PM, we utilized the ratiometric pHluorin sensors, PM-CYTO and PM-APO^11^. In CC^A^309-expressing cells, apoplastic pH was increased (≈ pH 0.5), while cytoplasmic pH was reduced (≈ pH 0.54), suggesting that the electrochemical potential across the PM is disturbed in the progress of CC^A^309-induced cell death (Fig. 2a, Extended Data Fig. 6). However, the pH change was moderate in PMA3 co-expressing cells (Fig. 2b), implying that CC^A^309 triggers alkalization of extracellular space by inhibition of PMAs. The early responses during HR are known to be followed by the activation of PM-bound ion channels and the assembly of an NADPH oxidase complex for the generation of reactive oxygen species^12^. Moreover, the elevation of cytosolic calcium places upstream of the oxidative burst in NLR-mediated cell death^13^. Thus, we monitored the influx of Ca^2+^ in cytosol and change of pH in apoplast during CC^A^309-induced cell death in *N. benthamiana* using cytosolic Ca^2+^ sensor, YC3.6 and PM-APO, respectively^14^. In the repeated experiments, we observed the elevation of cytosolic calcium ions at 6∼8 hrs post-induction of CC^A^309 expression which is significantly later than the elevation of pH at 2∼4 hrs in the apoplast (Fig. 2c, Extended Data Fig. 7). As a verification, artificial alkalization of the apoplast also triggered cytosolic Ca^2+^ accumulation (Fig. 2d). Furthermore, we observed the depolarization of PM by transient expression of CC^A^309 using DiBAC4 as an indicator^15^ (Fig. 2e). These results indicate that CC^A^309 triggers the alkalization of extracellular space and membrane depolarization via inhibition of PMAs leading to the influx of Ca^2+^ into the cytosol.

**Fig. 2.**
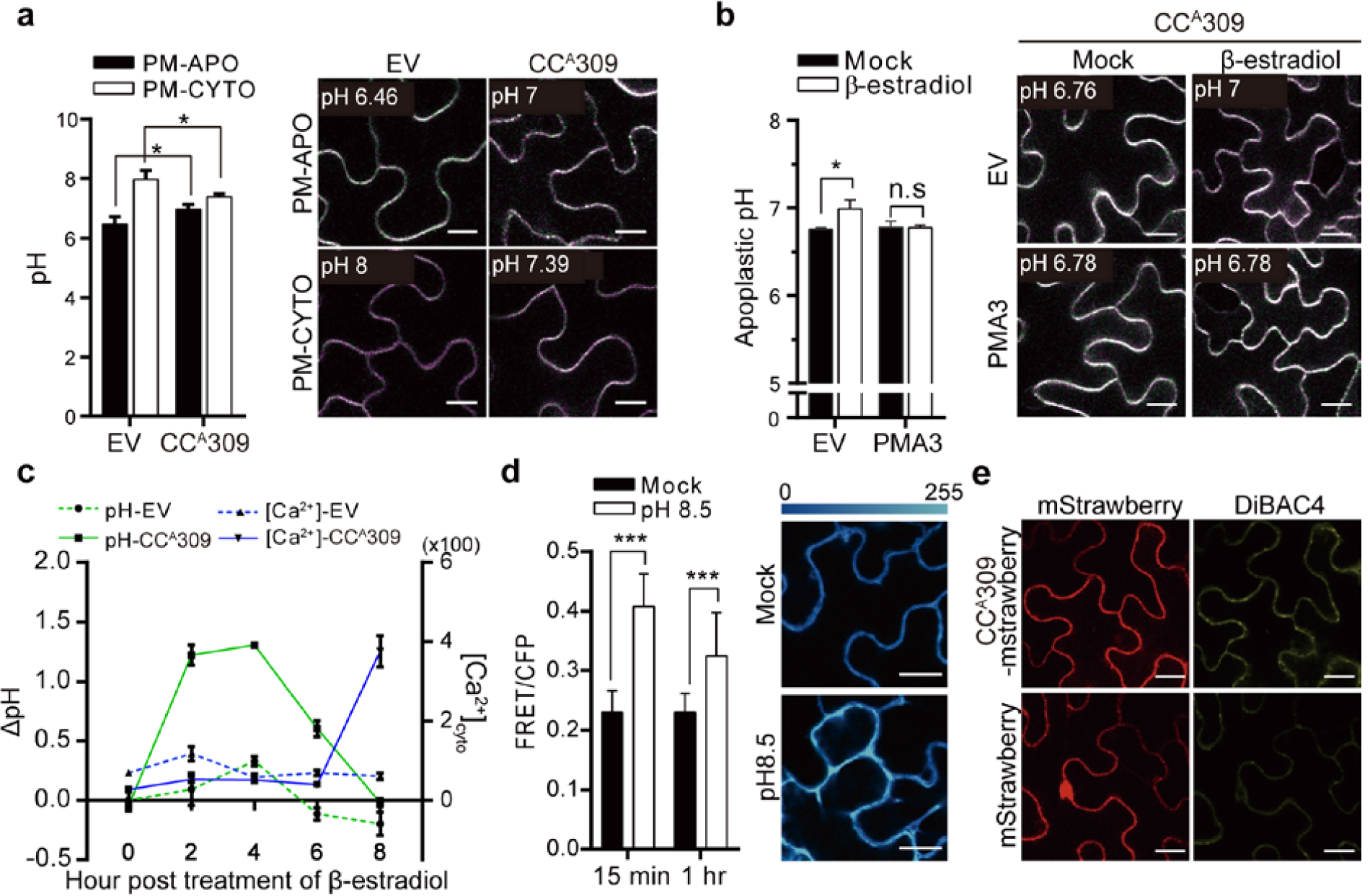
CC^A^309 triggers an extracellular alkalization leading to cytosolic Ca^2+^ influx. **a**, The elevated apoplastic pH and decreased cytoplasmic pH during CC^A^309-induced cell death in *N. benthamiana*. Beta-estradiol inducible *XVE:CC*^*A*^*309* or EV were co-expressed with apoplastic (PM-APO) or cytoplasmic (PM-CYTO) pH sensors. Bar = 15 μm. Data are represented as mean ± SD by triplicated independent experiments. Asterisk denotes a significant difference (t-test, *P < 0.05). **b**, The compromised CC^A^309-induced apoplastic alkalization by PMA3. PM-APO was co-expressed with EV or *35S:PMA3-FLAG*. The pH values (left) and representative confocal images (right) were taken 4 hr after induction. Data are represented as mean ± SD by triplicated independent experiments. Asterisk denotes a significant difference (t-test, *P < 0.05). Bar=15 μm. **c**, The change of apoplastic pH and cytoplasmic Ca^2+^ accumulation. PM-APO or cytosolic Ca^2+^sensor YC3.6 was co-expressed with EV or *XVE:CC*^*A*^*309*. Apoplastic pH and cytosolic Ca^2+^ concentration was measured at indicated time points. **d**, Induced cytosolic Ca^2+^ accumulation by apoplastic alkalization in YC3.6 infiltrated plant. 10 mM HEPES pH 7.0 (Mock) or pH 8.5 was infiltrated at 2 dpi. The Ca^2+^ concentration (left) and representative confocal images (right) were taken at 15 min and 1 hpi. Bar = 15 μm. **e**, CC^A^309-induced depolarization of plasma membrane (right top). The *35S:CC*^*A*^*309-mStrawberry* or *35S:mStrawberry* was expressed and stained with DiBAC4 at 8 hour post induction and photographed by confocal microscope. Bar = 20 μm. All experiment performed with 4-week-old *N. benthamiana*. Data are represented as mean ± SD (N = 10) by triplicated independent experiments. Asterisk denotes a significant difference (t-test, ***P < 0.001).

The PMA protein is composed of three major cytosolic domains, including a central catalytic and a C-terminal regulatory domain and is activated by phosphorylation on penultimate threonine allowing for the binding of the activator protein, 14-3-3 resulting in relieve of the autoinhibition state^16,17^. We found that CC^A^309 interacts with both central and C-terminal domains of PMA but a stronger co-IP signal was observed with the C-terminal domain (Fig. 3a).

**Fig. 3.**
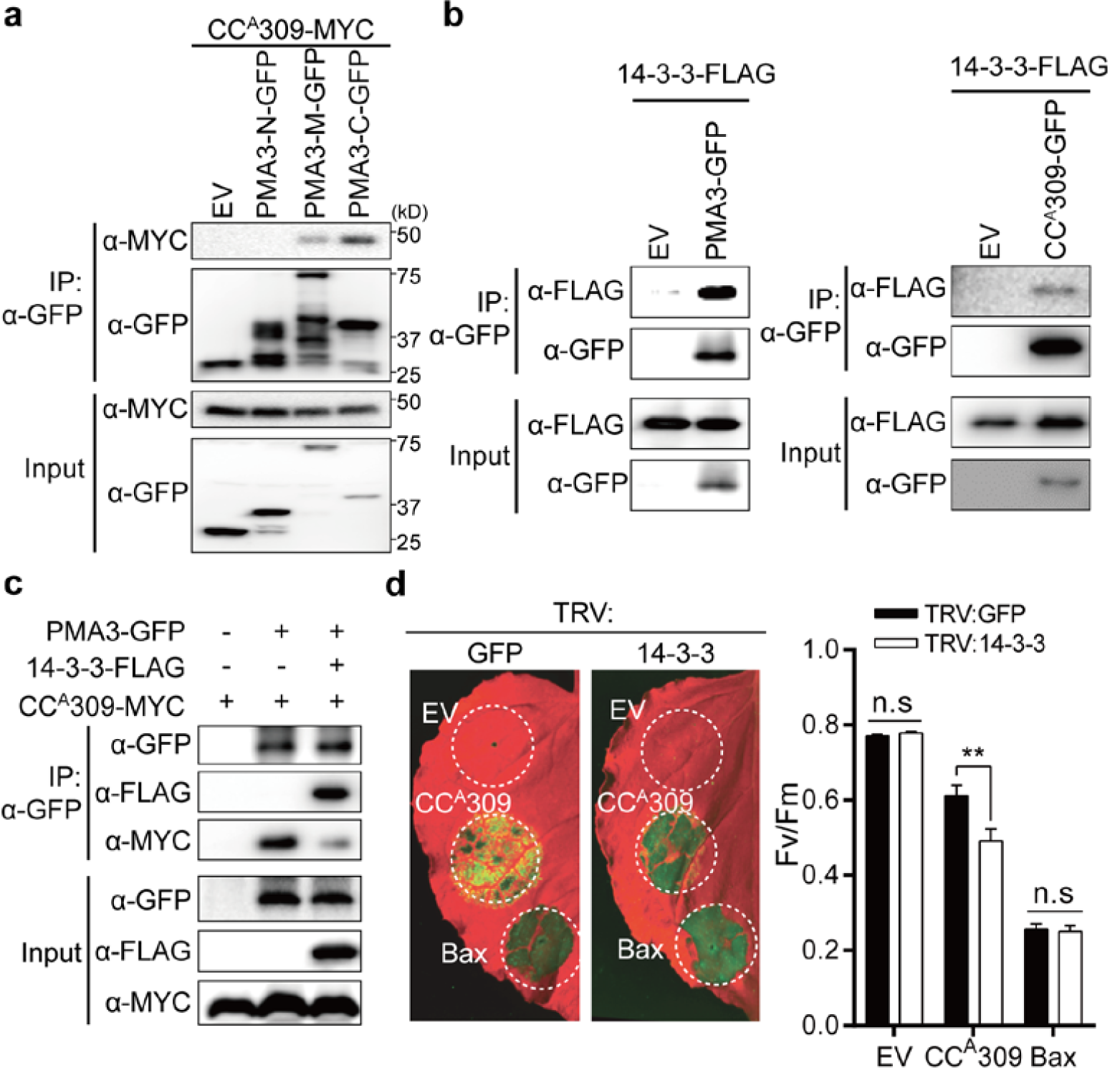
14-3-3 negatively regulates CC^A^309-induced cell death. **a**, CC^A^309 interacts with cytosolic central domain (PMA3-M) and C-terminal domain (PMA3-C), but not N-terminal domain (PMA3-N) of PMA3. *XVE:CC*^*A*^*309-MYC* was co-expressed with EV, *35S:PMA3-N-GFP, 35S:PMA3-M-GFP*, or *35S:PMA3-C-GFP*. Proteins extracts were immunoprecipitated with α-GFP (IP:α-GFP) and immunoblotted with α-MYC or α-GFP (top two panel). Protein inputs are shown with immunoblotting before IP (bottom two panels). **b**, CC^A^309, PMA3, and 14-3-3 form complex. *35S:14-3-3-FLAG* was co-expressed with EV and *35S:PMA3-GFP* (left panel) or *35S:CC*^*A*^*309-GFP* (right panel). Proteins extracts were immunoprecipitated with α-GFP (IP:α-GFP) and immunoblotted with α-FLAG or α-GFP (top two panel). Protein inputs are shown with immunoblotting before IP (bottom two panels). **c**, 14-3-3 inhibited the association between PMA3 and CC^A^309. PMA3-GFP and/or CC^A^309-MYC and/or EV were co-expressed with or without 14-3-3-FLAG. The co-IP was carried out with an α-GFP and analyzed by western blot with α-GFP, α-FLAG, or α-MYC (left top three panels). Protein inputs are shown with immunoblotting before IP (left bottom three panels). **d**, The enhanced *CC*^*A*^*309*-induced cell death in *14-3-3*-silenced plants. Three-week-old *N. benthamiana* silenced by *TRV:GFP* or *TRV:14-3-3* were agro-infiltrated with EV, *35S:CC*^*A*^*309-FLAG*, or *35S:Bax*. The leaves were photographed at 5 dpi and the degree of cell death is depicted as quantum yield (Fv/Fm). The Data are shown as mean ± SE (N > 12). Significance were determined using t-test, with asterisks denoting statistically significant differences. **P<0.01.

Because 14-3-3 is associated with both PMA3 and CC^A^309 in our co-IP experiment (Fig. 3b), the roles of 14-3-3 in CC^A^309-induced cell death were further investigated. Interestingly, co-expression of 14-3-3 and CC^A^309 compromised the association between CC^A^309 and PMA3 (Fig. 3c), suggesting that 14-3-3 may have an inhibitory effect on the interaction between PMA3 and CC^A^309. Consistent with co-IP data, significantly enhanced cell death was observed in *14-3-3*-silenced plants (Fig. 3d, Extended Data Fig. 8). Next, we determined whether PMA activity indeed affects CC domain-induced cell death. Fusicoccin, an irreversible activator of PMA by inhibiting dissociation of PMA/14-3-3, was used for further experiments^18^. The leaves expressing CC domains were infiltrated with fusicoccin. Surprisingly, only cell death induced by PM-localized and PMA3-associated CC domains (CC^A^309, CC^A^Pvr4, and CC^NbZAR1^) was inhibited by fusicoccin without deteriorative effect on protein expression, but not cytosol-localized CC^R3a^ (Fig. 4a,b,c, Extended Data Fig. 9a). To expand our understanding of PMA on resistance protein (R protein)-mediated cell death, functionally characterized NLRs were subjected to test fusicoccin effect. Full-length NLRs were expressed with or without its cognate effector in *N. benthamiana* and followed by fusicoccin treatment. As results, cell death mediated by PM-localized CNLs (Prf, HRT, Pvr4, RPS2, and NbZAR1^D481V^) but not non-PM-localized CNLs (R3a and Rx) and non-PM-localized TNLs (N, SNC1 and RPS4) was compromised by fusicoccin (Fig. 4d, Extended Data Fig. 9b). Moreover, full-length Pvr4 and NbZAR1 interact with PMA3, suggesting that PMA is a pivotal regulator of cell death in PM-associated R proteins (Extended Data Fig. 10)^19,20,21^.

**Fig. 4.**
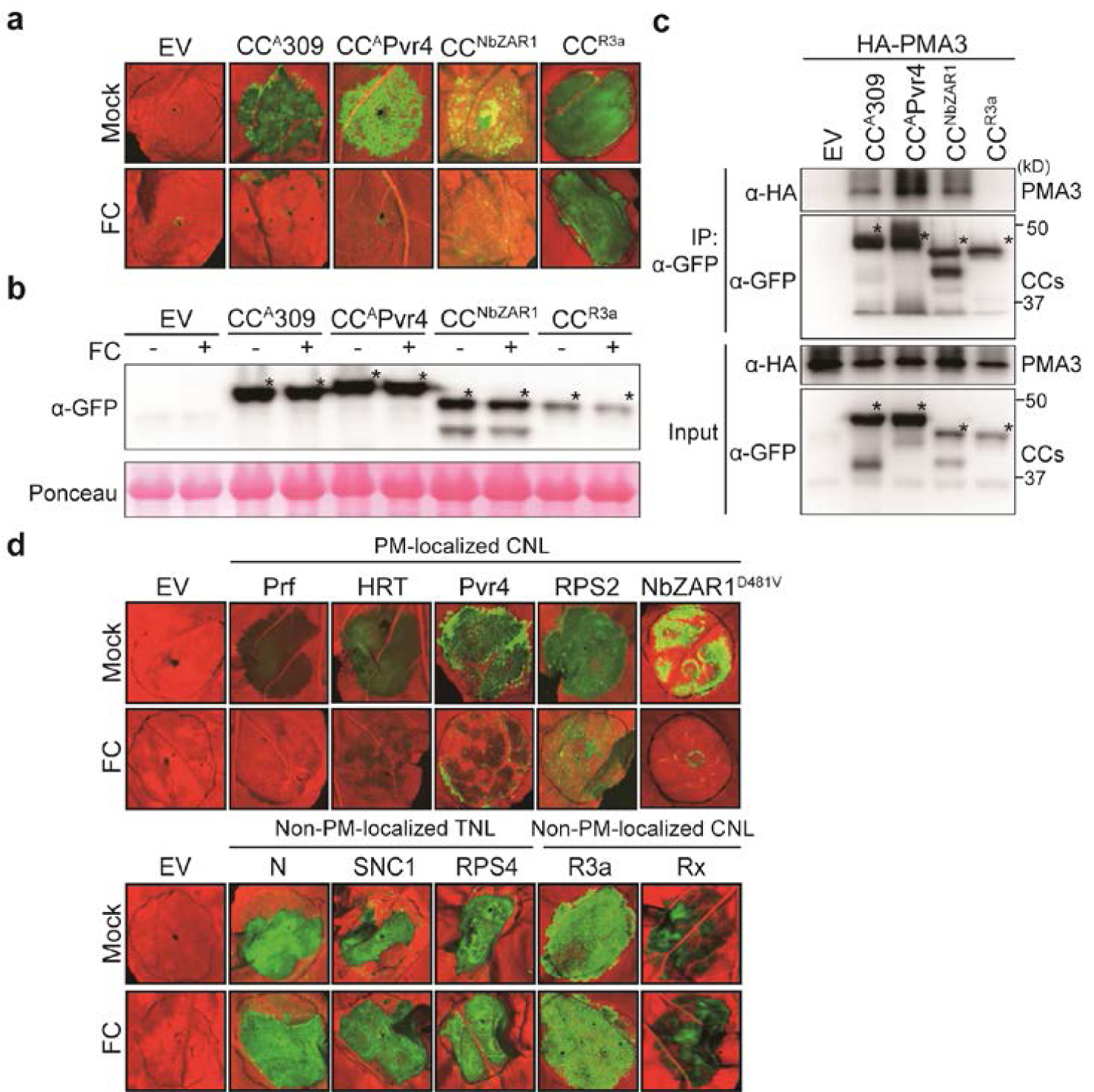
The cell death induced by PM-localized CNLs is inhibited by PMA activator. **a**, Fusicoccin (FC) inhibited cell death induced by CC domain of ANL309, Pvr4, NbZAR1, but not R3a. *Agrobacterium* carrying indicated construct were infiltrated in *N. benthamiana* then followed by infiltration of 1μM fusicoccin at 16 hpi. Leaves were photographed at 3 dpi. **b**, FC did not affect the protein expression. The proteins were detected by immunoblotting with α-GFP. Ponceau staining of RBC is loading control (bottom). **c**, The CC domains showing fusicoccin-dependent cell death inhibition are associated with PMA3. *35S:HA-PMA3* was co-expressed with CC domains. Proteins extracts were immunoprecipitated with α-GFP (IP:α-GFP) and immunoblotted with α-HA. Protein inputs are shown with immunoblotting before IP. Asterisks indicate the expected protein bands. **d**, FC inhibited cell death mediated by PM-localized CNL-type R proteins, but not TNLs or other organelle-targeting CNLs. Fusicoccin was treated in the same way as (**a)** at the infiltrated region by *Agrobacterium* carrying R genes and cognate Avr genes.

PM is the site of recognizing environmental cues and generating the secondary signals by modifications of PM-bounded channels and transporters. Although it is well-studied that the perception of pathogen results in the exchange of ion and solute across the PM as well as generation of reactive oxygen species (ROS), how activated NLR provokes cell death in immune response was obscure^22,23^. Our data clearly indicate that CNLs are associated with and inhibit PMA, resulting in alkalization of extracellular space leading to membrane depolarization. We propose PMA as a primary target of membrane-associated CNLs to facilitate both the development of cell death by disturbing homeostasis of membrane potential and the generation of secondary signals such as Ca^2+^ influx and ROS burst to trigger the defense response and HR cell death in plants (Extended Data Fig. 11).

## Methods

### Plant materials and growth conditions

*Nicotiana benthamiana* were grown in walk-in growth chamber at 25 °C with a 16 h/8 h (day/night) cycle. Four-weak-old plants were used in *Agrobacterium*-mediated transient expressions.

### Transient gene expression

For *in planta* transient overexpression, *Agrobacterium tumefaciens* strain GV3101 cells carrying the desired binary vector clones were grown overnight at 28 °C in LB media with appropriate antibiotics. The grown cells were harvested by centrifugation at 3,000 rpm for 10 min and resuspended into the infiltration buffer (10 mM MES (2-[N-morpholino] ethanesulfonic acid), 10 mM MgCl_2_ and 150 μM acetosyringone, pH5.6). Resuspended cell suspension were incubated for 1∼3 hr at room temperature and pressure-infiltrated with a 1 mL needless syringe into *N. benthamiana* leaves. To visualize cell death, the leaves were photograph after 3∼5 days after infiltration using Cy5 and Cy3 channels in Azure 400 (Azure biosystem).

### Constructs

Clones for epitope tagged proteins were generated using ligation-independent cloning method^24^. The inserts for *NbRIN4* (Niben101Scf03479g05049.1), *Nb14-3-3* (Niben101Scf02367g04001.1), *NbPMA1* (Niben101Scf00593g01002.1), *NbPMA3* (Niben101Scf07395g00031.1), *NbPMA4* (Niben101Scf03979g02010.1) were amplified from *N. benthamiana* cDNA and cloned into the pCAMBIA2300-LIC vectors (Cauliflower mosaic virus 35S promoter with N-term 6xHA, C-term eGFP or C-term 3xFLAG tag). For expression of fragments of PMA3, PMA3-N (residue 1-64), PMA3-M (residue 305-650) and PMA3-C (residue 846-956) were cloned into pCAMBIA2300-C-eGFP-LIC vector. For inducible expression, CC^A^309 (nucleotide 1-420) was cloned into XVE-DC-6xMYC vector by gateway cloning (Invitrogen)^25^. To generate single gene-silenced plants, the gene-specific fragment of *PMA3* (nucleotide 308-474) or 14-3-3 (nucleotide 54-338) was cloned into TRV2-LIC vector. For co-silencing of *PMA*s (PMA1/2/3/4), the fragments of *PMA1* (nucleotide 4-204), *PMA2* (nucleotide 202-401), *PMA3* (nucleotide 402-601), and *PMA4* (nucleotide 599-798) were fused by overlap PCR and cloned into TRV2-LIC vector. The CC domain of NbZAR1 (nucleotide 1-420), Pvr4 (nucleotide 1-495) or R3a (nucleotide 1-480) was fused to eGFP epitope by cloning to pCAMBIA2300-C-eGFP-LIC vector. The primers used for cloning are listed in Extended Data Table 2.

### Virus-induced gene silencing (VIGS)

VIGS was performed described by Liu et al^26^. Two-week-old *N. benthamiana* plants were used in VIGS. *A. tumefaciens* strain GV3101 harboring TRV1 and TRV2 with corresponding fragments from desired genes were grown overnight at 28 °C in LB media. Harvested cells were resuspended into infiltration buffer (10 mM MES, 10 mM MgCl_2_ and 150 μM acetosyringone, pH5.6) to a final O.D_600_ of 0.3. Resuspended cell suspension was incubated for 1∼3 hr at room temperature. Cell suspension of TRV1 and TRV2 were mixed in a 1:1 ratio before infiltration. At two weeks, the upper leaves were used for further experiments. To estimate the efficiency of silencing, transcripts level of genes was validated by quantitative RT-PCR. Total RNA was isolated from leaves of silenced plants using TRI reagent (MRC), and cDNA was synthesized using SuperScript II Reverse Transcriptase (Invitrogen). qRT-PCR was performed using a CFX96 Touch Real-Time PCR Detection Systems (Bio-rad) with SsoAdvanced Universal IT SYBR Green Supermix (Bio-rad). *Elongation factor-1 α* (*EF-1a*) was used as an internal standard. Gene-specific primers used for expression analysis are listed in Extended Data Table 2.

### Co-immunoprecipitation

*Agrobacterium*-mediated transient expression in *Nicotiana benthamiana* was performed as described with some modifications^27^. Briefly, *Agrobacterium* GV3101 (OD_600_=0.5) carrying different vectors tagged with HA, FLAG, GFP or MYC was syringe-infiltrated into 4-week-old *N. benthamiana* leaves. The infiltrated leaves were collected at 36 hpi for co-IP. The GFP tagged proteins were immunoprecipitated with 10 µl of α-GFP agarose beads (MBL) in 500 µl co-IP buffer (150 mM NaCl, 50 mM Tris-HCl, pH7.5, 2 mM EDTA, 10 mM DTT, 0.2% Triton X-100, 1:100 cOmplete protease inhibitor cocktail (Roche). A small aliquot of samples in co-IP buffer was used for input control before adding α-GFP agarose beads. The co-IP samples were gently rotated for 3 hrs at 4°C. The beads were collected and washed more than five times with washing buffer (500 mM NaCl, 25 mM Tris-HCl, pH7.5, 1 mM EDTA and 0.15 % NP-40. The samples were analyzed by immunoblot with an appropriate antibody.

### Bimolecular fluorescence complementation

pCAMBIA2300-LIC vector was modified for cloning BiFC construct to contain YN (residues 1-155 of YFP) or YC (residues 156 to the stop codon of YFP). NbRIN4 and CC^A^309 were fused with YN at their C-terminal in the pCAMBIA2300-LIC-YN vector. NbPMA3 was fused with YC at its C-terminal in the pCAMBIA2300-LIC-YC vector. NbRIN4-YN or CC^A^309-YN was transiently expressed with NbPMA3-YC and p19 silencing suppressor by agro-infiltration at a 1:1:1 ratio in *N. benthamiana* leaves. Then, treated with 2 mM LaCl_3_ at 16 hpi to inhibit CC^A^309-induced cell death. Imaging was performed using a confocal microscope (Leica SP8 X) at 2 dpi. To observe the BiFC fluorescence and chlorophyll autofluorescence, 514 and 633 nm white light laser were excited, respectively, and 525-580 and 650-720 nm emission wavelengths were used. Images were processed using LAS X software.

### Quantification of intensity of cell death

The intensity of cell death was quantified by measuring chlorophyll fluorescence using a closed FluorCam (Photon Systems Instruments) and Fluorcam 7.0 software. The detached leaves were exposed to the super pulse in the closed chamber and minimum fluorescence (F0), maximum fluorescence (Fm) and maximum quantum yield of photosystem II (Fv/Fm) parameters were determined by default F_-_ v/Fm protocol of system. The Fv/Fm value of empty vector-infiltrated leaves was around 0.77.

### Phylogenetic tree analysis

Amino acid sequences of PMAs in *Arabidopsis thaliana, N. benthamiana* and *N. plumbaginifolia* were aligned using Muscle^28^, and the alignment was used to construct a phylogenetic tree with the Maximum Likelihood method using MEGA7^29^. Evolutionary distances were computed using the JTT matrix-based method with bootstrap test (500 replicates).

### Fusicoccin treatment

Fusicoccin (1 mM, Sigma-Aldrich F0537) stock solution was prepared in ethanol and diluted in distilled water to 1μM. After 12-16 hpi, 1 μM fusicoccin or mock (0.1% ethanol) were infiltrated into the leaf region that has been infiltrated with *Agrobacteria* for transient expression of indicated genes. To determine if fusicoccin inhibits R protein-mediated cell death, *Pto/AvrPto* (for activation of Prf), *HRT/TCV-CP, Pvr4/PepMoV-NIb*, full-length of *RPS2, NbZAR1*^*D481V*^ (autoactive form of *NbZAR1*), *N/p50*, full-length of *SNC1*, full-length of *RPS4, R3a/Avr3a* and *Rx/PVX-CP* were expressed in *N. benthamiana*, then 1 μM fusicoccin was treated at 16 hr after infiltration.

### Confocal microscopy

To measure pH on either side of the adjacent plasma membrane, two retiometirc pHluorin sensors, PM-APO and PM-CYTO were received from Martiniere et al.^11^. *N. benthamiana* leaves was infiltrated with a mixture of *Agrobacteria* harboring pH sensor and XVE:CC^A^309. Two days after infiltration, 10 μM β-estradiol solution was sprayed on *Agrobacterium*-infiltrated region to activate CC^A^309 expression. Confocal microscopic observation and quantification of intensity of fluorescence signals were followed as described with modifications. Observations were performed with a Leica SP8 X microscope, using a 20X water objective with the same WLL laser at 20% of 476 nm and at 20% of 496 nm output. Emission light was at a pinhole of 1 Airy between 505 and 550 nm. For DiBAC_4_(3) imaging, leaf discs were incubated in DiBAC_4_(3) (10 μM, Invitrogen B438) for 30 min and we used the excitation line at 488 nm and recovered fluorescence signal between 505 and 560 nm. Calibration curves were obtained *in situ* with the purified proteins from *E. coli* diluted in buffers of various pH levels. Yellow Cameleon 3.6 (YC3.6) was expressed in the same way as pH sensors. The 405 nm of a diode laser was used to excite CFP and FRET signal was collected in 525-535 nm. To quantify the intensity of the fluorescence signal, images were analyzed using Image J software. After the subtraction of background noise, the average mean gray value was calculated for each channel and ratio images were generated.

### Statistics

Graphs were generated by PRISM 7 (graphPad) and figure legends indicate type of statistical test used. Error bars generally represent standard deviation or standard error.

## Acknowledgements

We are thankful to Dr. Nadine Paris who provided pHluorin sensors and Dr. Cecile Segonzac for critical reading of the manuscript. This work was supported by the National Research Foundation of Korea (NRF) grant funded by the Korea government (MSIT) (No. 2018R1A5A1023599 (SRC) and 2018R1A2A1A05019892).

## Author contributions

D.C. conceived the project; H.L., H.M. Y.S., and D.C. designed the experiments. H.L., H.M., Y.S., J.L., S.O., S.L., J.K., S.J., H.K., H.P., S.K. performed the experiments. H.L., H.M., Y.S., and D.C. wrote the manuscript.

## Competing interest declaration

Authors declare no competing interests.

## Supplementary Data

**Extended Data Fig.1.**
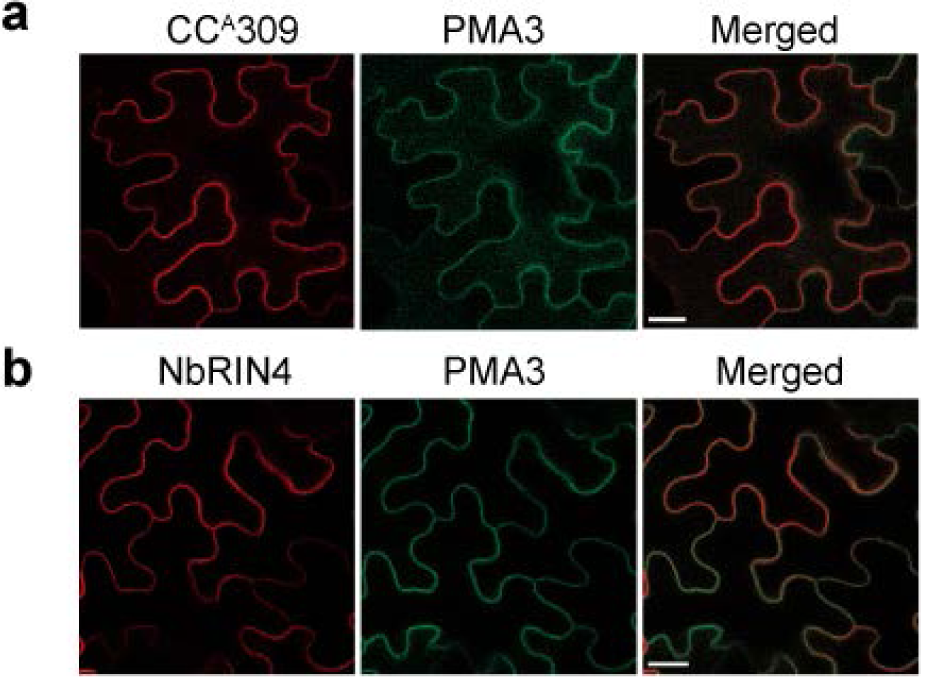
CC^A^309, NbRIN4 and PMA3 are co-localized at plasma membrane. The subcellular localization of the CC^A^309, NbRIN4, and PMA3 were determined in *N. benthamiana* epidermal cells. CC^A^309, PMA3, and NbRIN4 localized in plasma membrane. *Agrobacterium* carrying CC^A^309-mstrawberry (***a***) or NbRIN4-mstrawberry (***b***) were mixed with PMA3-eGFP and infiltrated in 4 week-old *N. benthamiana*. At 2 dpi, the fluorescence signals were observed by confocal microscopy. The images were obtained by combination of Z-stack overlays. Bar=20 μm.

**Extended data Fig. 2.**
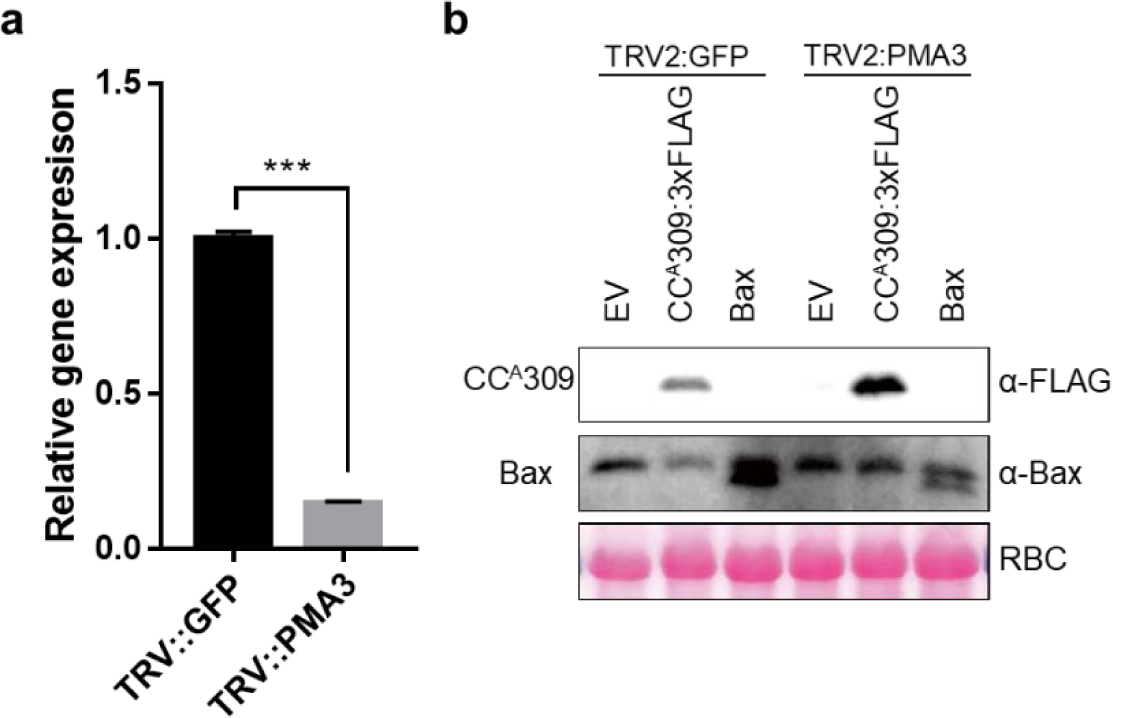
Virus-induced gene silencing of *PMA3* and protein levels of overexpressed genes in *PMA3*-silenced plants. **a**, Decreased transcript of *PMA3* in *PMA3*-silenced plant. The transcript level of *PMA3* was measured by quantitative RT-PCR at 2 weeks after VIGS. The mean values (± SD) for transcript levels were normalized to that of *N. benthamiana EF1-α*. Transcript levels *PMA3* in *GFP*-silenced plants were set to 1. Error bars represent the mean of three biological replicates. Asterisks denote significant differences at ***P < 0.001 as determined by t-test. **b**, Protein accumulation in *GFP*- or *PMA3*-silenced plants was confirmed by immunoblot analysis. Equal protein loading was confirmed by Ponceau staining of membrane. Asterisks indicate the expected protein bands.

**Extended Data Fig. 3.**
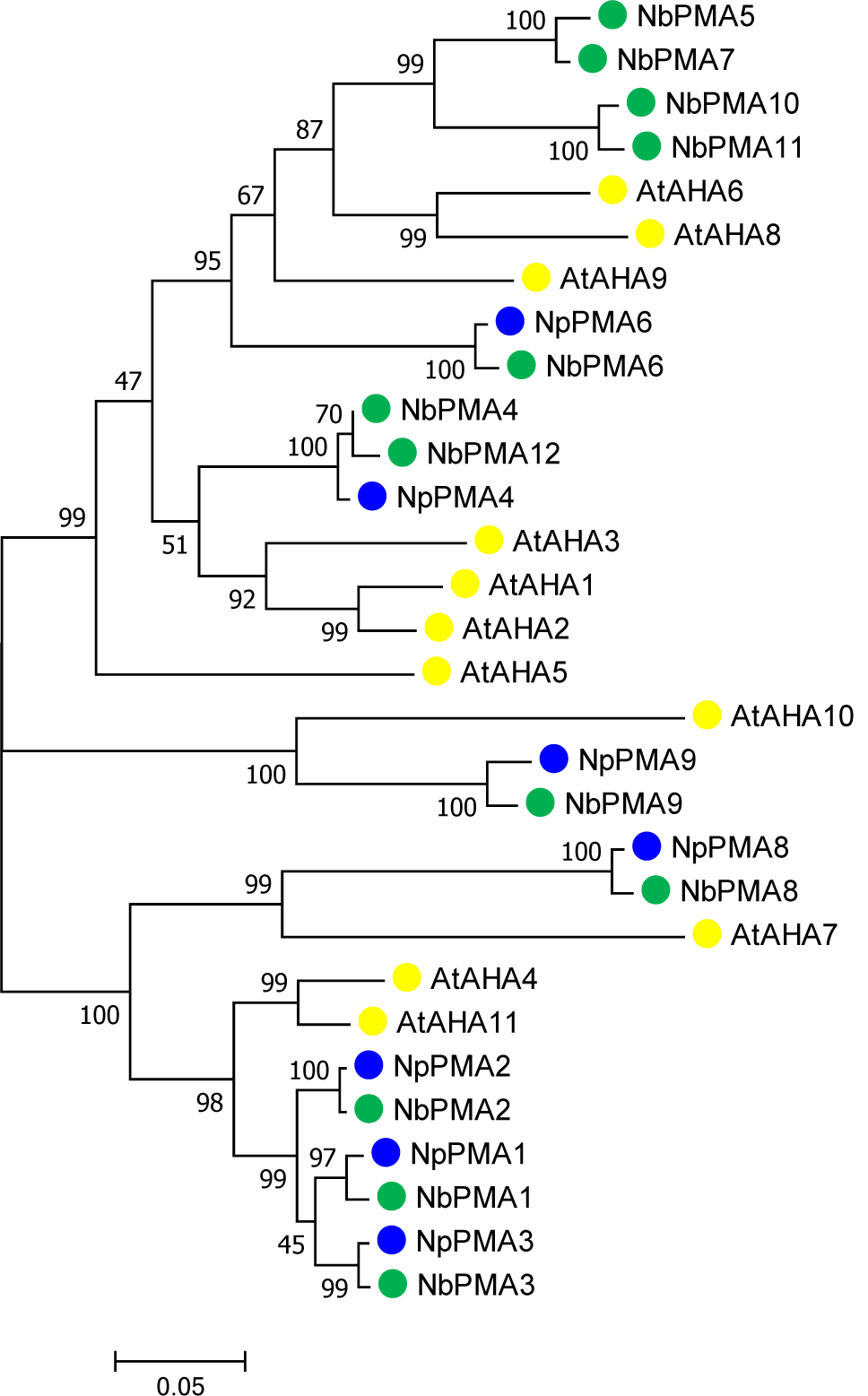
Phylogenetic analysis of PMAs in *Arabidopsis, N*. *benthamiana* and *N*. *plumbaginifolia*. Amino acid sequences of 11 PMAs in *Arabidopsis thaliana*, 12 PMAs in *N. benthamiana* and 9 PMAs *N. plumbaginifolia* were aligned for construction of phylogenetic tree. Yellow; *Arabidopsis* PMAs; blue, *N. plumbaginifolia*; green, *N. benthamiana*. Phylogenetic analysis was conducted by a maximum likelihood method with 500 replications of bootsrapping in MEGA7.

**Extended Data Fig 4.**
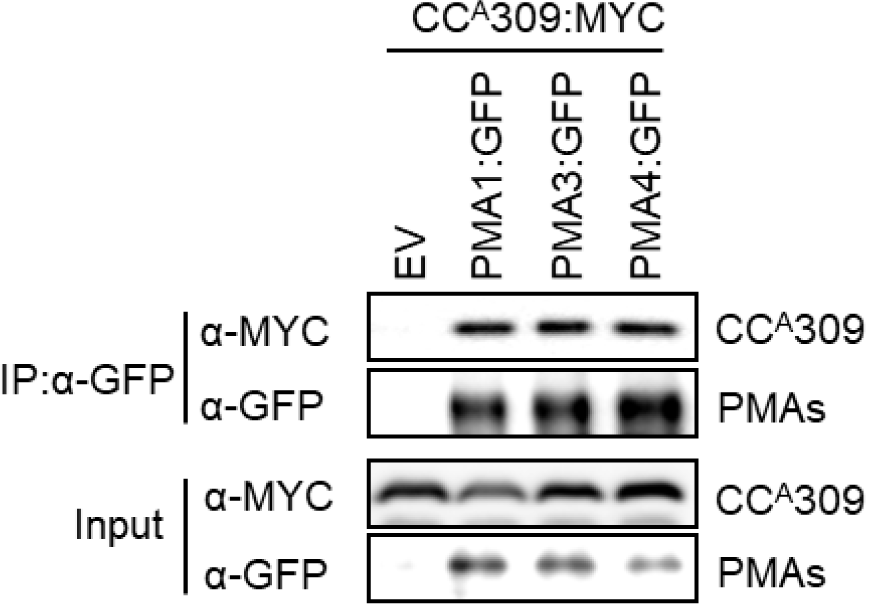
PMA1 and PMA4 as well as PMA3 interact with CC^A^309. NbPMA1, NbPMA3, and NbPMA4 interact with CC^A^309. NbPMA1-, NbPMA3- and PMA4-GFP were co-expressed with CC^A^309-MYC in *N. benthamiana*. Protein extracts were immunoprecipitaed with GFP antibody (IP: α-GFP) and immunoblotted with α-GFP or α-MYC (top two panels). Protein inputs are shown with immunoblotting before immunoprecipitation (bottom two panels).

**Extended Data Fig 5.**
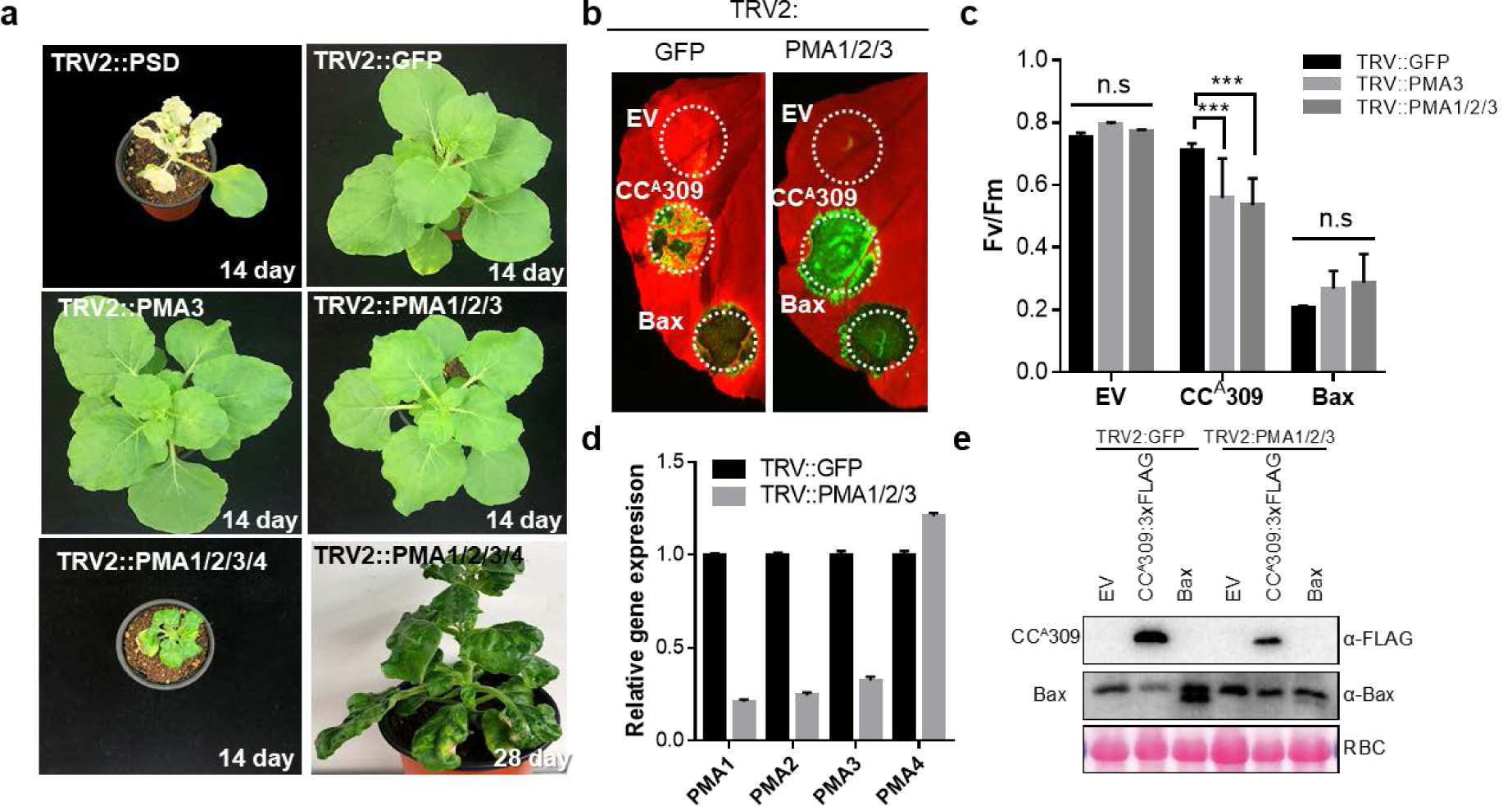
CC^A^309-induced cell death is enhanced in *PMAs*-silenced plants. **a**, The phenotypes of *NbPMA3-, NbPMA1/2/3-* or *NbPMA1/2/3/4*-silenced plants. Abnormal growth phenotypes were observed in *PMA1/2/3/4*-silenced plant. The photobleached leaves of *phytoene desaturase* (*PDS*)-silenced plant represent the effective gene silencing. *GFP*-silenced plant was used as control for VIGS. Pictures were taken at 14 or 28 days after VIGS. **b**, The CC^A^309-induced cell death is enhanced in TRV2:*PMA1/2/3* in compared to TRV2:*GFP* plants. The plants at 10 days after VIGS were used to be infiltrated with *Agrobacteria* harboring EV, CC^A^309-3xFLAG and Bax. The leaves were photographed at 3 dpi. **c**, The data represents the degree of cell death using the quantum yield (Fv/Fm). Data are represented as mean ± SD (n = 9). **d**, The transcripts level of *NbPMA1, NbPMA2, NbPMA3* and *NbPMA4* in *PMA1/2/3*-silenced plants were measured to estimate the silencing efficiency by qRT-PCR. The mean values (± SD) for transcript levels were normalized to that of *N. benthamiana EF1-α*. **e**, Protein accumulation of CC^A^309 and Bax in *GFP*- or *NbPMA1/2/3*-silenced plants was confirmed by immunoblot analysis. Protein loading was confirmed by Ponceau staining of membrane. Asterisks indicate the expected protein bands.

**Extended Data Fig 6.**
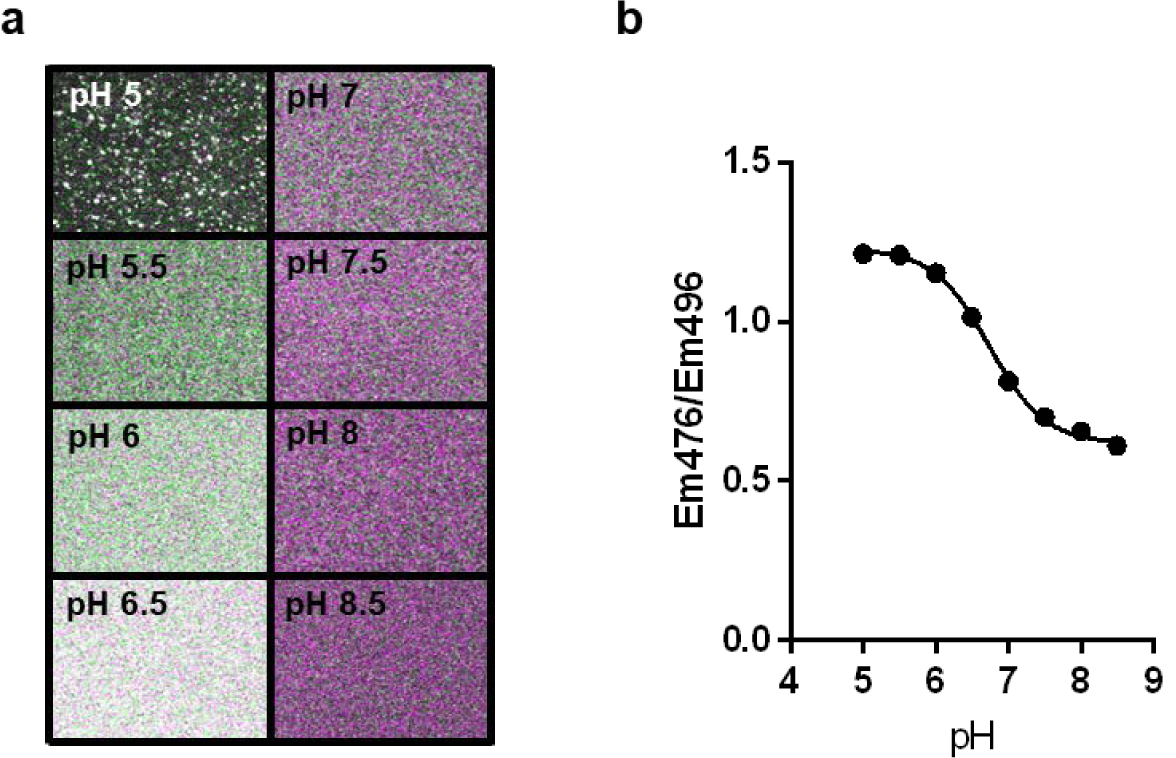
pHluorin response to external pH and calibration curve. **a**, Pseudocolored fluorescence images of pHluorin in buffers with different pH values. The purified pHluorin proteins from *E. coli* liquid culture were diluted in 50 mM MES-KOH buffers at pH5, 5.5, 6, 6.5 or in 50 mM HEPES-KOH at pH 7, 7.5, 8, 8.5. The buffers containing the pHluorin dropped onto slide glass and illuminated at 497 nm and 496 nm using a confocal microscope. **b**, Calibration curve was obtained from (**a**) to define absolute pH values.

**Extended Data Fig 7.**
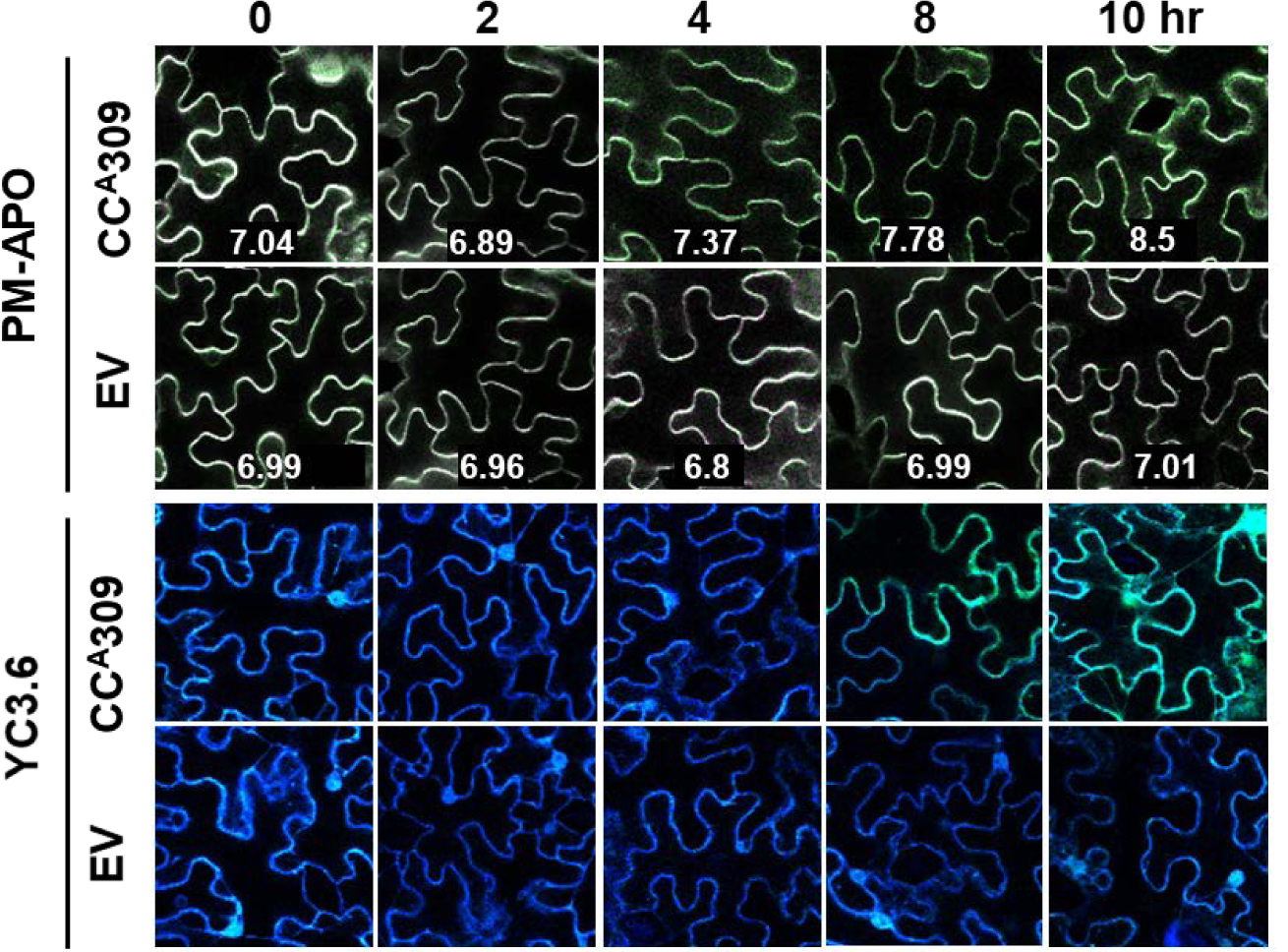
The earlier alkalizations in apoplast then cytosolic Ca^2+^ accumulation during CC^A^309 expression. Pseudocolored fluorescence images of PM-APO and YC3.6 were obtained from transiently expressed leaves infiltrated with EV or inducible CC^A^309 and PM-APO or YC3.6 in *N. benthamiana*. respectively. Pictures were taken at various time points after spray of 10 μM of β-estratiol solution to induce CC^A^309.

**Extended Data Fig 8.**
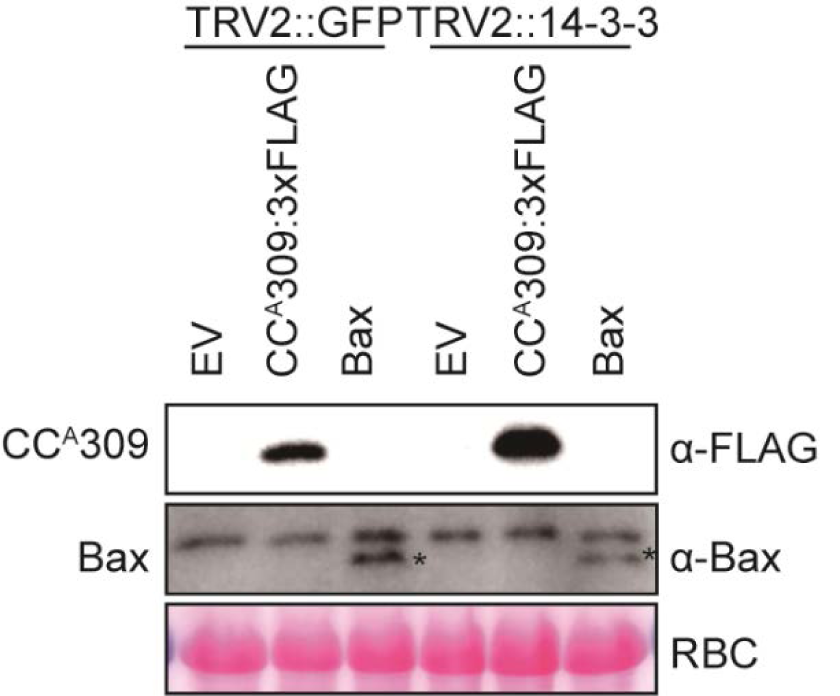
Protein accumulation of CC^A^309 and Bax in *GFP*- or *Nb14-3-3*-silenced plants. Protein accumulation of CC^A^309 and Bax in *GFP*- or *14-3-3*-silenced plants was confirmed by immunoblot analysis with α-FLAG or α-Bax antibody (top two panels). Equal protein loading was confirmed by Ponceau staining of membrane (bottom panel). Asterisks indicate the expected protein bands.

**Extended Data Fig 9.**
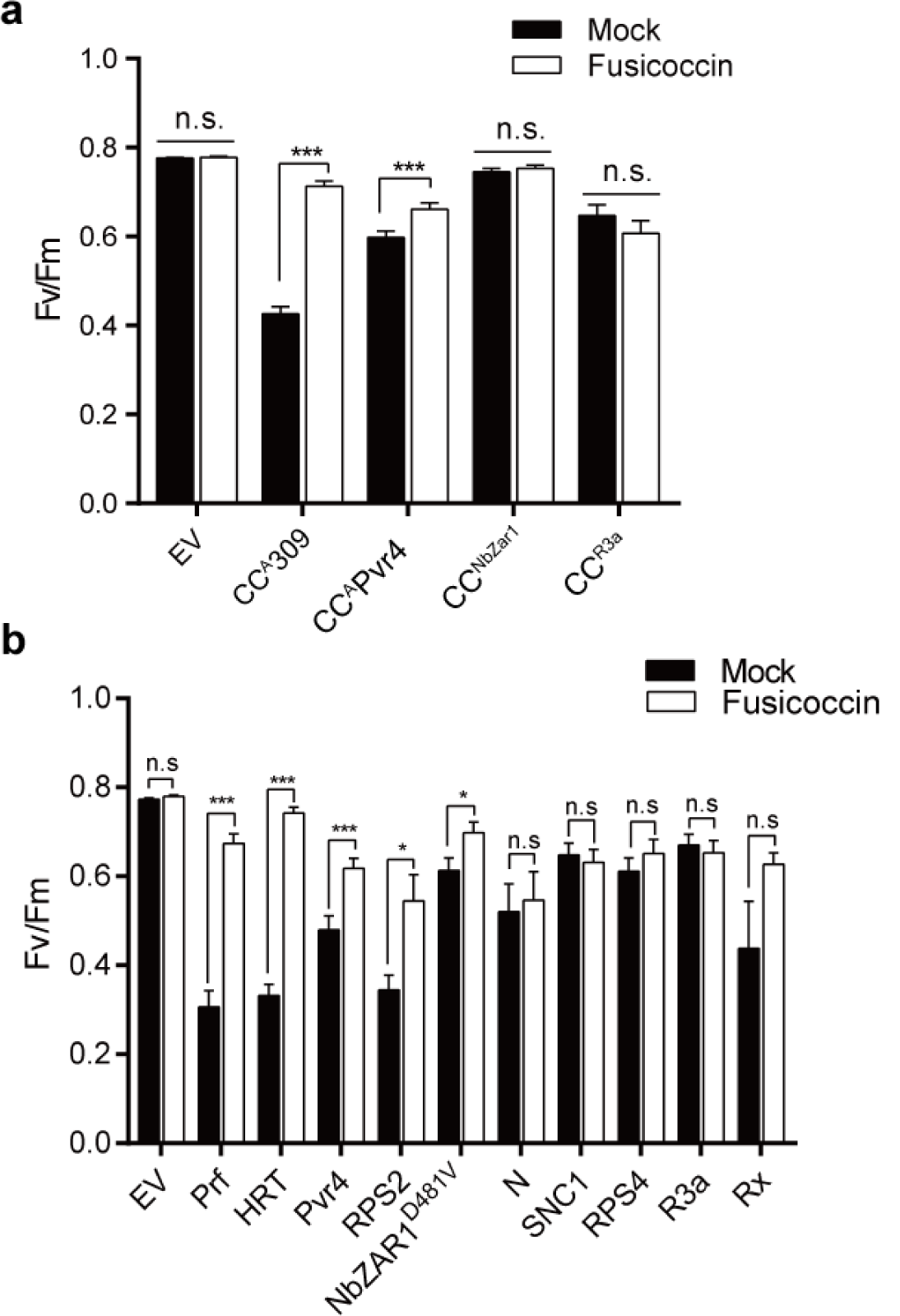
The cell death induced by CC domains or R proteins was inhibited by treatment of fusicoccin. ***a, b***, The degree of cell death in Fig. 4a, d is depicted by quantification of quantum yield (Fv/Fm). Data are represented as mean ± standard error (n = 5∼25) by triplicated independent experiments. Significance was determined using t-test. *<0.05, **p<0.01, ***P<0.001.

**Extended Data Fig 10.**
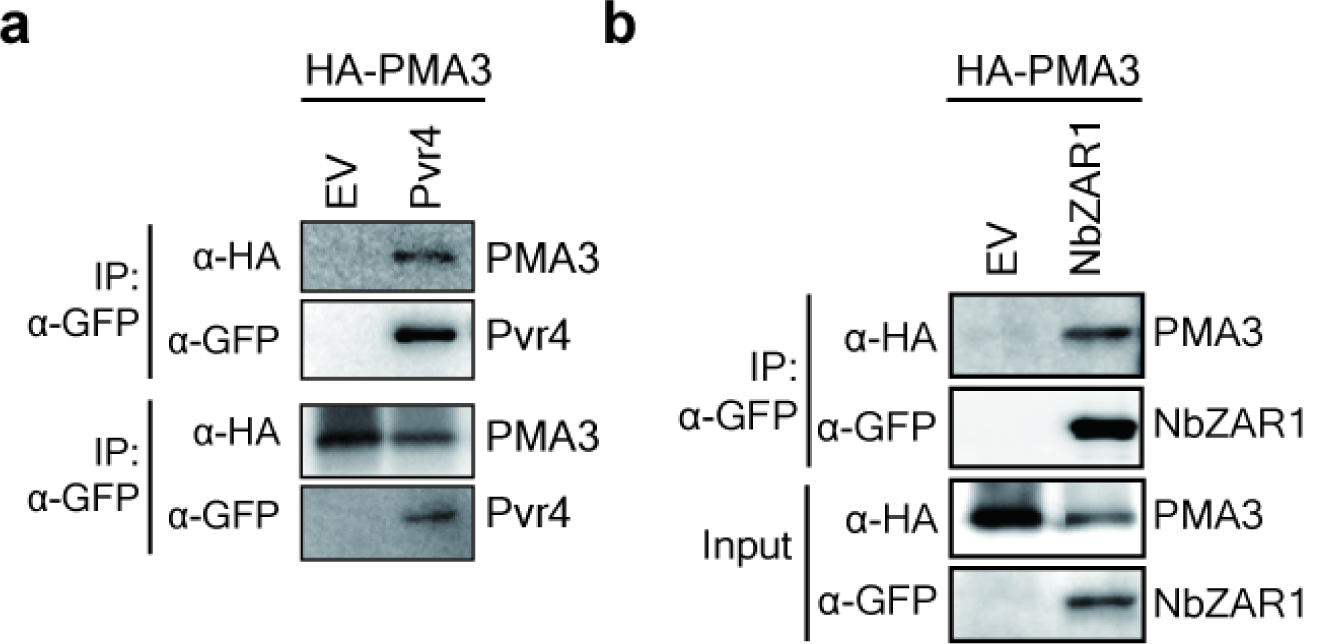
Full-length R proteins, Pvr4 and NbZAR1 also interact with PMA3. *35S:HA-PMA3* was co-expressed with EV and *35S:Pvr4-GFP a* or *35S:NbZAR1-GFP b*. Proteins extracts were immunoprecipitated with α-GFP (IP:α-GFP) and immunoblotted with α-HA or α-GFP (top two panel). Protein inputs are shown with immunoblotting before IP (bottom two panels).

**Extended Data Fig 11.**
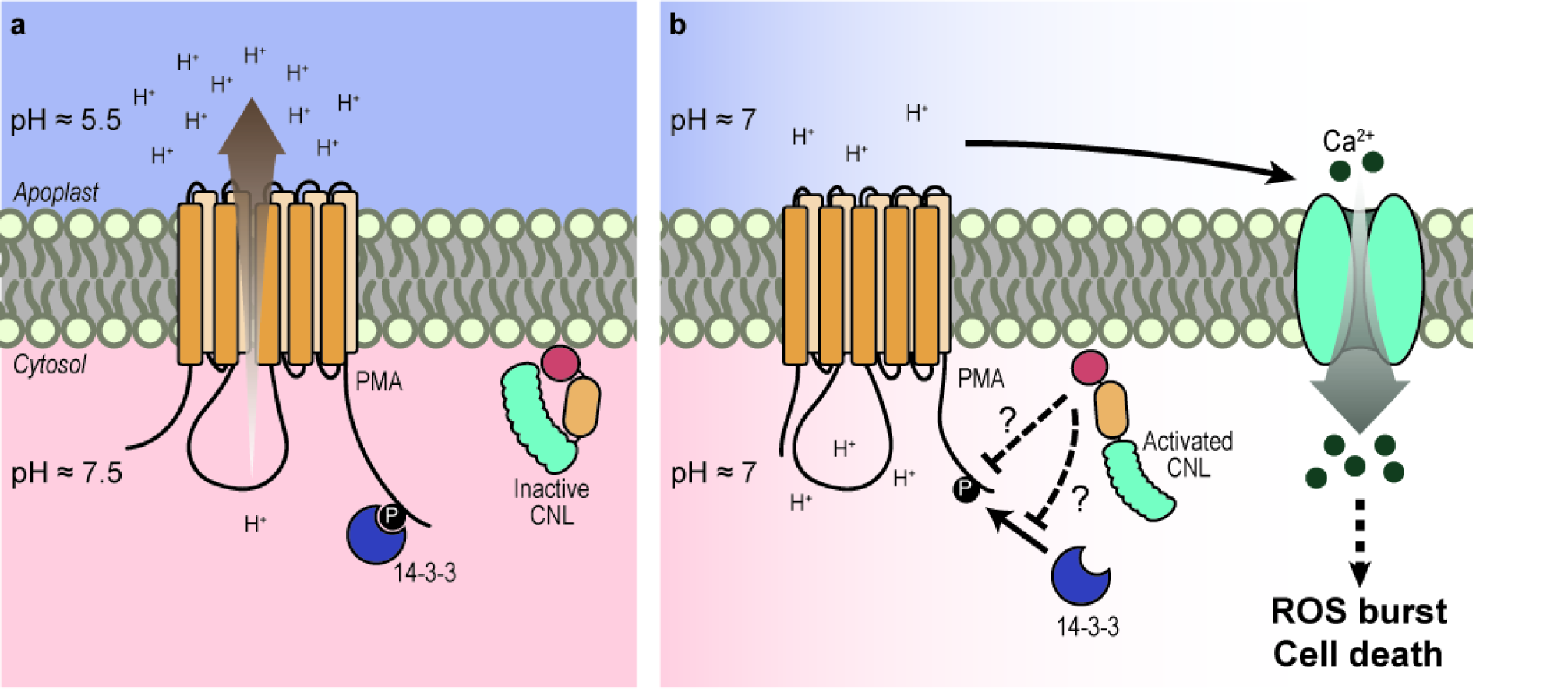
Proposed mechanism of plasma membrane-associated CNL-mediated cell death. **a**, The rest state of polarized plasma membrane. **b**, PM depolarization triggered by activated CNLs. The change of PM potential might facilitate a series of PM-integrated defense responses such as Ca^2+^ influx and ROS burst, and consequently leads to cell death.

**Extended table 1.** List of the CC^A^309 interactor candidates identified by mass spectrometry

| Gene description | Gene ID | # of peptides |  | Co-IP |
| --- | --- | --- | --- | --- |
|  |  | 35S:GFP-3xFLAG | 35S:CCA309-3xFLAG |  |
| CCA309 |  | 0 | 223 |  |
| GFP |  | 1352 | 0 |  |
| ATP synthase epsilon chain | Niben101Scf01399g00012.1 | 2 | 30 | + |
| Acetyl-CoA carboxylase | Niben101Scf04077g03006.1 | 0 | 26 | - |
| Cell division protein FtsZ homolog 1 | Niben101Scf03107g01011.1 | 0 | 24 | n.t |
| Chaperonin 60 | Niben101Scf02842g00014.1 | 0 | 21 | - |
| Glutathione S-transferase | Niben101Scf08250g03016.1 | 1 | 21 | n.t |
| EPSIN2 | Niben101Scf01066g01009.1 | 2 | 18 | n.t |
| Peptidyl-prolyl cis-trans isomerase FKBP16-3 | Niben101Scf06423g01016.1 | 0 | 18 | + |
| Cinnamoyl-CoA reductase 2-like | Niben101Scf09883g01010.1 | 0 | 16 | - |
| Acyl-CoA-binding protein | Niben101Scf02240g02008.1 | 0 | 13 | n.t |
| Thioredoxin superfamily protein | Niben101Scf10490g00011.1 | 0 | 12 | - |
| Thiamine pyrophosphate enzyme | Niben101Scf02538g06008.1 | 0 | 11 | - |
| Coiled coil domain containing protein | Niben101Scf10747g00018.1 | 0 | 10 | n.t |
| Chaperonin 60-2 | Niben101Scf29076g00011.1 | 0 | 10 | - |
| Macrophage migration inhibitory factor | Niben101Scf00302g02011.1 | 0 | 10 | - |
| Reactive Intermediate Deaminase A | Niben101Scf04339g07004.1 | 0 | 9 | + |
| ATP synthase delta chain | Niben101Scf15372g00003.1 | 0 | 8 | - |
| Translation elongation factor EF1B | Niben101Scf03350g00002.1 | 0 | 7 | - |
| Trigalactosyldiacylglycerol 2 | Niben101Scf07377g00001.1 | 0 | 7 | + |
| pre-mRNA-processing factor 39 | Niben101Scf11535g01033.1 | 0 | 7 | n.t |
| 2-methylene-furan-3-one reductase | Niben101Scf01155g00008.1 | 0 | 7 | - |
| Plasma membrane ATPase 3 | Niben101Scf07395g00031.1 | 0 | 6 | + |
| Peroxiredoxin | Niben101Scf03671g02013.1 | 0 | 6 | - |
| Superoxide dismutase | Niben101Scf04451g00026.1 | 0 | 5 | - |
| Tripeptidyl-peptidase 2 isoform | Niben101Scf00595g08041.1 | 0 | 5 | n.t |
| Light-harvesting complex II proteins | Niben101Scf06008g00011.1 | 0 | 4 | + |
| Zinc finger A20 and AN1 domain-containing protein | Niben101Scf05348g04001.1 | 0 | 3 | - |

**Extended table 2.**
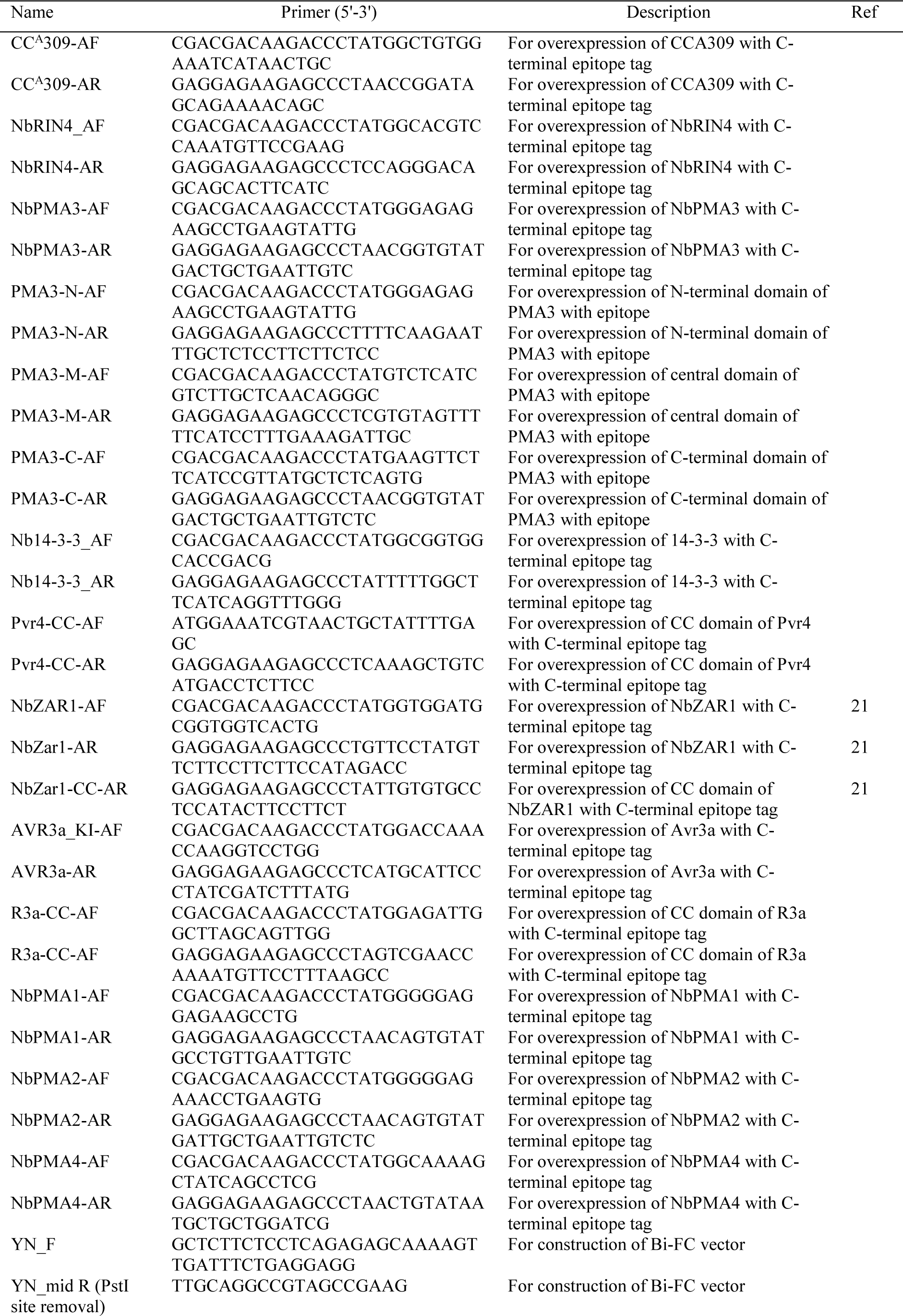
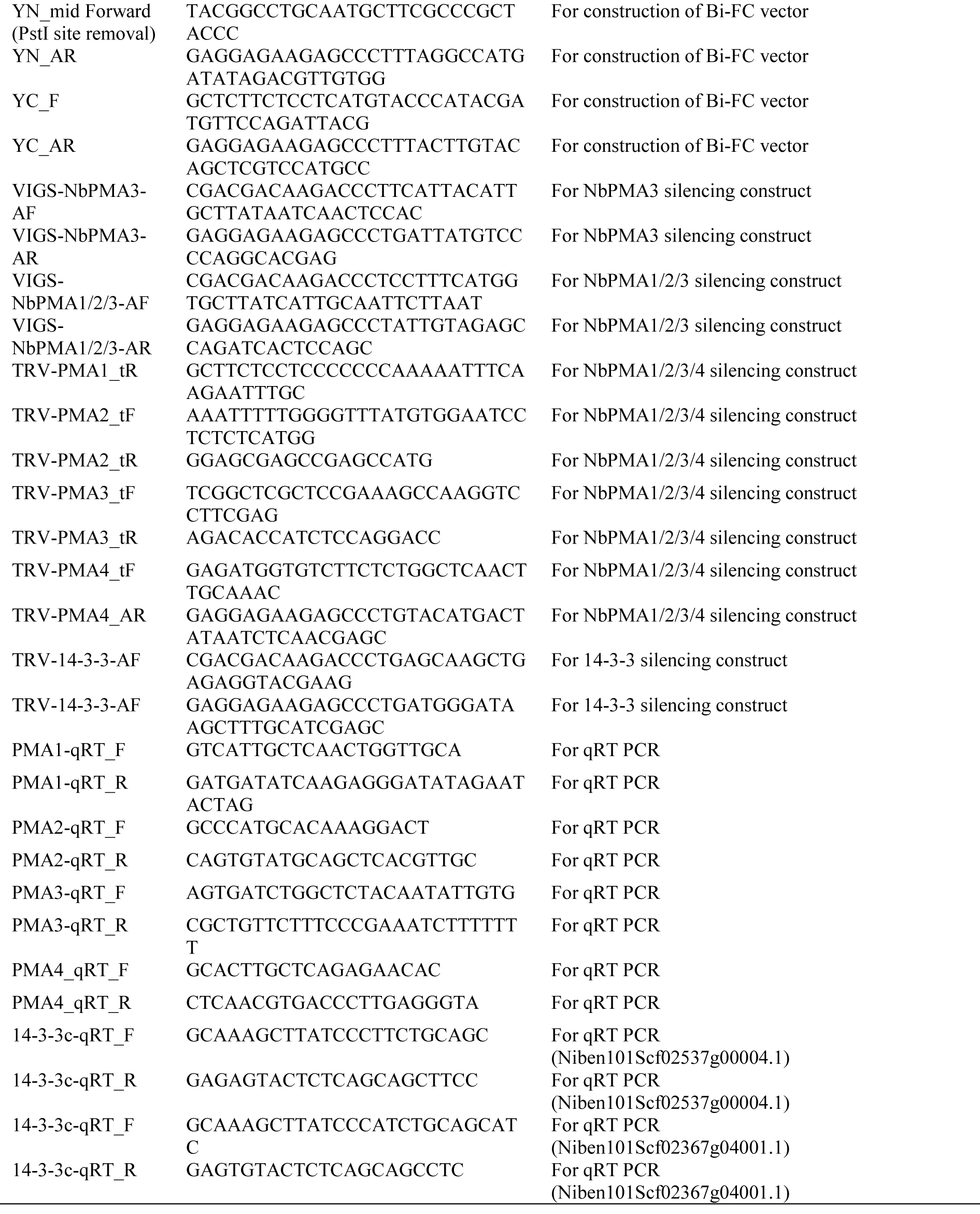
List of primers used in this study

## Notes

### Competing Interest Statement

The authors have declared no competing interest.

